# Can a history of crop rotations improve the prediction of soil organic carbon in the Andes? integrating machine learning multi-annual crop classification as a proxy of soil management

**DOI:** 10.64898/2025.12.06.692759

**Authors:** Marcelo Bueno, Hildo Loayza, Johan Ninanya, Javier Rinza, Percy Briceño, Luis Silva, Carlos Mestanza, Ronal Otiniano, Jan Kreuze, David A. Ramírez

**Affiliations:** Programa de posgrado en Cambio Climático, Universidad Nacional de San Antonio Abad del Cusco, Cusco, Peru; International Potato Center (CIP), Headquarters, P.O. Box 1558, Lima 15024, Peru; Departamento Académico de Suelos, Universidad Nacional Agraria La Molina, Lima, Peru; Universidad de Huánuco, Huánuco, Peru; Asociación Pataz, Jr. Independencia 263, Trujillo, Peru

## Abstract

Soil organic carbon (SOC) is a crucial component related to various processes that ensure soil health and function. Its modeling is vital for assessing and monitoring soil degradation caused by the potential impact of agricultural activities. This study aimed to model SOC in the Northern highlands of Peru, characterized by a high amount of SOC, which is being affected by crop expansion. Crop rotation (CR) was incorporated into a modeling exercise using remote sensing data, fieldwork, and farmer surveys. A multi-year classification model with seven cropland classes was developed using data collected from 534 fields across 2022-2024, including 189 soil samples. Each cropland field was represented as a polygon delineating its boundaries and indicating its dominant crop cover. Time series of multispectral Sentinel-2 Level-2A Top of Canopy imagery were used to derive phenological features—such as the timing of maximum canopy cover and the length of the growing period—based on Normalized Difference Vegetation Index (NDVI) time series. A Random Forest classifier was used as the baseline model. The cropland classification model demonstrated strong overall performance, with F_1_ scores ranging from 0.81 to 0.98 across the different classes. The model performed well for lupin and pasture but scored lower for beans and potatoes. Predictions of cropland classes from 2019 to 2022 were created, resulting in frequency layers that represent crop rotations. Four feature configurations were evaluated: (i) including all features as a benchmark, (ii) excluding climatology, (iii) excluding crop rotation history, and (iv) excluding soil properties. Configurations including all features and excluding crop rotation history showed the highest performance (*R*^2^ = 0.63), while those excluding climatology or soil properties performed worse (*R*^2^ ≈ 0.52–0.53). Although soil features were the most important, fallow frequency emerged as the most critical predictor of SOC in crop rotations. When soil data were excluded, fallow frequency, combined with climatic features, explained over half of the SOC variability. The findings emphasize the importance of incorporating CR into SOC mapping efforts.

## Introduction

To meet the growing global demand for food, agricultural crop production has intensified in recent decades. This intensification threatens soil quality and increases atmospheric greenhouse gas levels [1]. Soil organic carbon (SOC) is widely recognized as a crucial component of soil quality in agricultural systems, playing a significant role in sustaining crop production and ensuring food security. Furthermore, sequestering SOC is considered a promising method to offset greenhouse gas emissions and facilitate carbon neutrality [2]. However, intensive conventional agricultural practices have hurt soil quality, particularly by reducing SOC. To address these issues, it is crucial to enhance SOC accumulation in agroecosystems through the adoption of sustainable agricultural management practices. Crop rotation (CR) is one such practice in regenerative agriculture [3], where a series of crops are grown sequentially over time on the same croplands [4, 5]. The adoption of CR can enhance agricultural production and soil fertility [6]. Additionally, CR significantly impacts SOC dynamics, as crop species diversity in CR systems directly affects the quantity and quality of residue-derived carbon input into the soil [7]. However, the degree and variability of SOC accumulation in response to CR, along with the underlying factors driving this response, remain uncertain [8–10]. Due to the interactions among various influential factors—such as climate, soil properties, and agronomic practices—there is considerable uncertainty in predicting SOC content responses to CR based on individual variables (e.g., air temperature, precipitation, soil texture, and crop types) [8]. Information about CR can be gathered from annual crop inventories, government maps, surveys, and earth observation (EO) technologies. Using satellite image time series has emerged as a promising approach for crop type classification in remote sensing. It captures spectral-temporal profiles from multispectral images taken throughout the growing season, reflecting the phenology of crops and their seasonal variations. This approach enhances classification accuracy by extracting unique spectral-temporal features for various crops [11–13]. However, a notable gap remains in knowledge regarding the integration of soil management into agricultural remote sensing frameworks.

Andean farming systems are typically mixed, combining livestock with a high level of inter- and intra-specific crop diversity. These systems primarily rely on crops such as potato (*Solanum* spp.), oca (*Oxalis tuberosa*), olluco (*Ullucus tuberosus*), mashua (*Tropaeolum tuberosum*), maca (*Lepidium meyenii* ), and quinoa (*Chenopodium quinoa*), as well as older crops like barley (*Hordeum vulgare*) and oats (*Avena sativa*) [14]. Different cropping rotations in the Andes are often accompanied by sectoral fallows, known locally as “laymis,” within communal territories [15]. These fallow periods help restore soil nutrients [16]. However, due to socioeconomic factors and population growth, these regions’ fallow periods have been reduced [17]. The encroachment of crops into highland grasslands has increased in the Andes, primarily driven by global warming [18, 19]. This trend poses a significant threat to the SOC stocks preserved by these grasslands [20]; thus, urgent restoration activities are needed to recover these ecosystems [21]. To enhance the quantification of SOC and analyze the key activities that help preserve soil carbon, it is crucial to focus on the Andes, which holds great potential for carbon sequestration [18, 22]. This study aimed to assess the importance of crop rotation features in predicting SOC. Multi-year crop rotation covariates were developed to evaluate their contribution to SOC variability and to provide insights into cropping practices that support sustainable soil management. The specific objectives of the study were to: i) develop a phenological-based multi-year (2019–2023) cropland classification model integrating Sentinel-2 time series; ii) explore the potential of CR history features for SOC prediction, while also evaluating alternative feature sets—climatology and soil properties—to assess the relative importance of these feature groups. The working hypothesis was that including CR features would enhance SOC prediction in heterogeneous landscapes, such as croplands in the highland Andes, even when soil features are not available.

## Materials and methods

### Study area

The study was conducted in the highlands of the Chugay district (7*^◦^*46*^′^*56*^′′^*S, 77*^◦^*52*^′^*03*^′′^*W, 3371 m a.s.l.), La Libertad region of Peru, focusing on a study area of approximately 255 km^2^ (Fig 1A). This area is dominated by grasslands, where smallholder agriculture has been established, with potatoes as the main crop [23].

**Fig 1.**
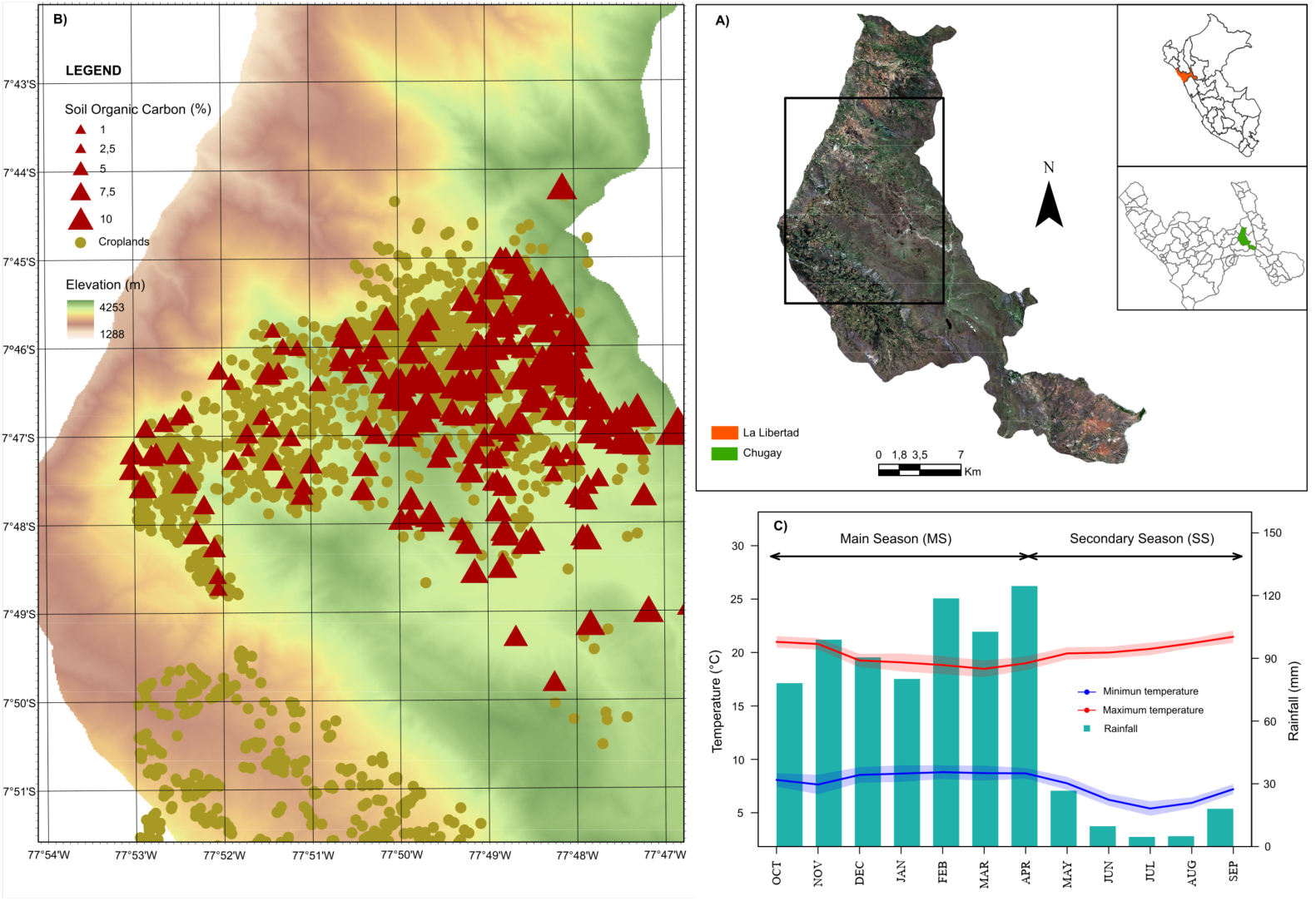
The study area. The study area (indicated by a rectangle over a Sentinel-2 scene from 17/08/2019) is situated in the Chugay District, La Libertad Department, Peru (A). Red triangles denote soil organic carbon sampling points, while green patches depict cropland fields in the study area (B). Monthly mean minimum (blue) and maximum (red) temperatures, and monthly accumulated precipitation (sky-blue bars) for the 1981–2016 period, derived from PISCO datasets (C).

Potatoes are typically grown under two rotation schemes: (i) fallow – potato – cereal (oat, barley, wheat) – legume (lupin, bean), and (ii) fallow – potato – cereal – fallow.

Fallow refers to arable land left uncultivated to recover from intensive cropping, typically covered by low-density *Rumex acetosella* weed. Potatoes are primarily cultivated during the rainy season, from October to April of the following year. To better understand the cropping dynamics in this region, it is important to distinguish between the main (MS) and secondary (SS) growing seasons. MS aligns with the onset of rains from late September to early October, during which potatoes are the predominant crop. SS may occur between May and September, although its timing is highly variable due to fluctuations in soil moisture availability. As a result, the MS spans two calendar years, while the SS falls within a single year, reflecting both inter-and intra-annual cropping phenology.

The climate is rainy and temperate with dry autumns and winters [24]. Monthly minimum and maximum temperatures range from 5.4 to 8.0 *^◦^*C and from 18.5 to 21.5 *^◦^*C, respectively, with an average annual rainfall of 757.7 mm, approximately 90% of which occurs during the potato growing season (Fig 1C), based on the 1981–2016 period from PISCO datasets [25].

According to the national soil map of Peru [26], the study area is predominantly characterized by Eutric Regosols–Eutric Cambisols (RGe–CMe code; i.e., young to moderately developed soils with slightly acidic to neutral pH and high surface organic matter), with smaller areas of Eutric Leptosols–Lithic outcrops (LPe–R code; i.e., shallow, somewhat acidic soils with coarse fragments near the surface) in the north and south. These soils are suitable for grazing or pastures, but present limitations for other crops due to edaphic constraints [27], as they are naturally acidic and lack carbonates despite their high organic matter content [28].

### Cropland labels and digitization of prediction masks

Cropland cover information of 189 fields was collected in 2022 and 2024 through field campaigns [29] and farmer surveys. Planting and harvesting dates for most fields in 2022 and 2024 were validated directly in the field through observation and surveys conducted with farmers. The coordinates of the central point of each field were obtained using a handheld GPS (Fig 1B). This was essential for aligning the crop monitoring points spatially with the farmers’ fields for soil sampling (Section 2.8) and subsequent remote sensing analysis (Section 2.5).

The field boundaries were determined using central point GPS coordinates and Planet imagery with a 1.5 m ground sample distance (GSD), acquired in August 2022 [30], ensuring synchronized field visits for crop monitoring. Each cropland was represented as a polygon delineating the field boundaries and its dominant crop type coverage (fallow lands were treated as an additional category) from 2022 to 2024, for both the MS and SS. To increase the number of samples for cropland classification, a dataset of 345 potato (*Solanum tuberosum* L.) fields containing only sowing and harvest information from the 2021 MS was included from the e-Agrology platform database [29]. All non-agricultural land covers (e.g., water bodies, forests, and urban areas) were omitted because SOC sampling was conducted exclusively in agricultural areas.

In addition to these 534 monitored cropland fields used for training and evaluation (Section 2.6), most of the croplands in the study area were identified and digitized, resulting in an additional 1,843 cropland fields. The digitization process used high-resolution imagery [30] for precise delineation. Finally, this dataset aimed to create a comprehensive cropland and non-cropland mask, supporting the generalization of predictions for the crop classification model (Section 2.6).

### Cropland label processing

The cropping calendar in the study area is highly variable due to the region’s diverse climate conditions. Therefore, to support season-specific modeling, the Sentinel-2 (S2) imagery time series was grouped by season type into MS and SS, enabling separate analysis of interannual and intra-annual cropping periods. To address class imbalance, only the most frequent crop classes, representing 97% of the dataset, were kept while less frequent crops (i.e., quinoa—*Chenopodium quinoa*—and corn—*Zea mays*—) were dropped.

Additionally, fields that were too small (≤ 250 m^2^) or excessively narrow, as identified in the original database, were removed to minimize mixed pixels and facilitate classification following [31] recommendations. Finally, the croplands considered were: barley (*Hordeum vulgare* L.), fallow, oat (*Avena sativa* L.), pasture, lupin (*Lupinus mutabilis*), beans (*Vicia faba*), and potato (*Solanum tuberosum* L.) (S1 Fig). These cropland labels generally span several seasons within the same field, thus providing a reliable historical record.

In this study, an instance is defined as a uniquely labeled cropland field for a specific season (therefore, a single field may have multiple instances—for example, potato MS 2023, barley SS 2024, etc.). These instances served as the basis for subsequent S2 time series acquisition and multi-year Machine Learning (ML) modeling (Section 2.6).

### Sentinel-2 time series acquisition

A custom algorithm was developed to download Harmonized Level-2A surface reflectance time series for each cropland using the Google Earth Engine (GEE) Python API and geemap [32]. The data were retrieved from the COPERNICUS/S2 HARMONIZED ee.ImageCollection, which provides 10 m GSD visible and near-infrared (NIR) bands and 20 m GSD shortwave infrared (SWIR) bands. Scenes with *>*70% cloud cover were excluded using the Scene Classification Layer metadata, while residual cloudy pixels were subsequently masked using a cloud probability layer [33]. Data were downloaded from 2019 to 2024; however, for 2024, data were only collected up to September, before the start of a new seasonal cycle, which was not considered in this study. The S2 SWIR bands were resampled to 10 m GSD using the Python library Xarray version 2023.3.0 [34]. Subsequently, reflectance time series were smoothed using the Savitzky–Golay filter implemented in the signal R package [35].

Since planting and harvesting dates identified through field surveys were not always reliable or available, crop growth windows were estimated using a consistent criterion. The zonal medians of NDVI [36], derived from red and NIR reflectance (see S1 Table), and their temporal patterns were analyzed for each cropland field, using medians instead of means to reduce the influence of outliers, given that the median is more robust [37]. Because NDVI reflects vegetation vigor and biomass, higher values are typically observed during peak growth stages (e.g., flowering), while values near zero are indicative of planting and senescence dates [38]. Finally, plausible planting and harvesting dates were estimated through preliminary surveys and then refined using local knowledge of cropping calendars, rainfall onset (e.g., in Andean rainfed agriculture, planting dates are determined by rainfall patterns), and visual inspection of NDVI time series.

### Sentinel-2 time series cropland phenological analysis

Achieving crop separability during each growing season involved combining three types of features that were used for multi-year cropland type classification modeling (S1 Table): (i) phenological characteristics derived from NDVI time series, (ii) vegetation indices, and (iii) preliminary clustering of NDVI time series of cropland types using Dynamic Time Warping (DTW).

First, a Gaussian function was used to model crop seasonal phenology, fitting NDVI time series in a way that preserves the maximum amount of information while simultaneously reducing the dimensionality of the data [39], which was parameterized using the following equation:

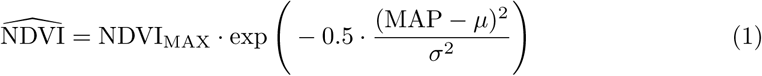

here NDVI_MAX_ is the NDVI value at peak canopy cover time, *µ* is the time to reach maximum canopy cover, and *σ* is the full width at half maximum of the fitted curve, related to the growing period length. Months After Planting (MAP) tracks crop development time. Only cropland fields with at least eight valid Sentinel-2 scenes were retained. The Gaussian function was fitted using the gslnls library in R [40] by non-linear least squares using the Marquardt method [41]. The resulting parameters—NDVI_MAX_*, µ*, and *σ*—were used as features for cropland classification.

To test for significant differences among cropland covers for the Gaussian parameters, the Kruskal–Wallis rank sum test was used, followed by Dunn’s post hoc test with Holm’s correction for multiple comparisons. The analyses were performed using the R packages stats [40] and DescTools [42]. Additionally, box plots were used to present the distribution of fitted Gaussian parameters by cropland type.

Second, the zonal medians from vegetation indices—bare soil index (BSI), specific leaf area vegetation index (SLAVI), normalized burn ratio 2 (NBR2), and normalized difference moisture index (NDMI) (see S1 Table)—were evaluated at the time of maximum canopy cover, i.e., closest to *µ* for each cropland type. Vegetation index calculations and zonal medians were performed in Python using Xarray version 2023.3.0.

Third, Dynamic Time Warping (DTW) clustering [43] was used to group similar NDVI growth patterns. DTW uses the Euclidean distance for similarity measurement between two temporal sequences. Given two time series sequences *U* = {*u*_1_*, u*_2_*, . . . , u_n_*} and *V* = {*v*_1_*, v*_2_*, . . . , v_m_*}, the base distance matrix *D*_base_ ∈ R*^n×m^* (where each element stores the Euclidean distances between points in the two sequences) is defined as:

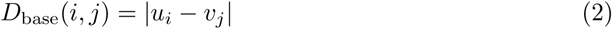

The DTW distance matrix *D* ∈ R*^n×m^* is obtained recursively as:

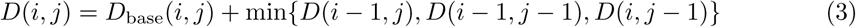

The lower the distance between two time series, the more similar they are. These distances were used to perform unsupervised clustering of cropland type temporal patterns using the dtwclust R library [44]. The distance parameter was set to dtw basic, and partitional clustering was applied with *k* = 7. Fuzzy clustering was also performed using the TSclust library [45], with clusters set to *k* = 7 in both cases to match the number of cropland types under study.

### Multi-year cropland type classification modeling

Given the sample size and label imbalance, a single multiannual model was trained using instances from 2021–2024, following the approach of [46] and [47]. Multiannual training helps capture time-independent features, averages out meteorological variations, and mitigates the impact of class imbalances by increasing the dataset size. The Random Forest (RF) algorithm [48] was used as the modeling framework due to its proven effectiveness for cropland classification with limited labeled data [13, 49].

To further address class imbalance in the dataset, data augmentation was applied using the themis library in R with the SMOTE–ENC algorithm [50], setting *k* = 5 and over ratio=1. Before augmentation, categorical features such as the DTW cluster groups were transformed into dummy variables using the fastDummies package.

Hyperparameter tuning was conducted using the tuneRanger package [51], which performs automatic model-based optimization [52] through the ranger package [53]. The tuned parameters included mtry (number of variables randomly sampled at each split), min.node.size (minimum number of samples in a leaf node), and sample.fraction (fraction of observations used for each tree), while the number of trees (num.trees) was fixed at 1000. The tuning process relied on out-of-bag accuracy as the performance criterion, allowing for faster yet robust estimation compared to cross-validation. Finally, with the optimal model configuration, the model was refitted using the optimized parameters on a 70% split of the complete dataset, following the recommendations of [54].

A confusion matrix [55] was used to summarize the classification results on the remaining 30% test set. Because the dataset was highly imbalanced, overall accuracy could be biased toward dominant cropland types (e.g., potato). To address this, the F1 score [56] was calculated for each class. Overall accuracy was also reported for comparison. Predictions were then performed for all cropland fields without an observed crop history.

A working example of the procedure suggested in this publication, including source code and example data, is provided on GitHub (https://github.com/kundun14/multi-annual-crop-classification-with-phenological-features).

### Multi-year cropland frequency as a rotation feature

A trained cropland classification model was used to enhance the soil management history, integrating the predicted cropland maps into a comprehensive crop rotation dataset. With the fine-tuned and validated model, crop type information from the 189 surveyed fields was extended to cover the 2019–2022 period. Integrating crop sequence information facilitated the incorporation of rotation history into our SOC modeling framework. Cropping frequency was subsequently derived from these extended sequences to quantify the prevalence of each crop category over the study period according to the following equation:

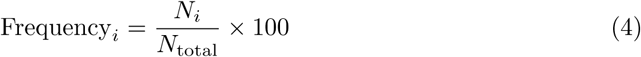

where Frequency*_i_* is the cropping frequency (%) for crop *i*, *N_i_* is the number of seasons in which crop *i* was detected in a given field, and *N*_total_ is the total number of seasons analyzed.

These frequency features reflect the dominance of various crop types over time and may help explain spatio-temporal trends in SOC [57]. This methodology resulted in seven frequency raster layers, one dedicated to each crop type.

### Soil sampling

In August 2022, 189 locations (Fig. 1B) were sampled using composite sampling of five points centered in the middle of the farmers’ fields, following the procedure of [21]. Soil samples were taken at 0–24 cm depth for physical-chemical analysis and at 15 cm for dry bulk density (Bd) analysis with the core method [58], which involved extracting two samples from the center of the croplands (0.15 and 0.30 m deep) in two volumetric cylinders with diameters of 50 mm and heights of 50.8 mm.

Soil samples were analyzed in 2023 at the Soil Laboratory of the La Molina National Agrarian University (Lima, Peru), following protocols described by [59]. The measured variables included pH (1:1 soil-water suspension), electrical conductivity (EC) (1:1 extract), available phosphorus (P) (Olsen method), available potassium (K) (buffered acetate extraction), soil organic carbon (SOC) using the Walkley & Black method, cation exchange capacity (CEC) determined with buffered ammonium acetate at pH 7.0, exchangeable cations—calcium (eCa), magnesium (eMg), potassium (eK), and sodium (eNa)—and exchangeable acidity (eAC) analyzed by atomic absorption spectrophotometry.

Particle size distribution (sand, silt, and clay) was determined using the Bouyoucos hydrometer method, and exchangeable acidity was also assessed. Additionally, the nitrogen isotopic composition (*δ*^15^N) of dried soil samples was analyzed at the Stable Isotope Facility, University of California, Davis [60].

### Features for soil organic carbon content prediction

The features used to predict SOC were classified into three groups (S2 Table): (i) soil physical-chemical properties, (ii) gridded historical climate, and (iii) multi-year cropland frequencies.

For the second set of features, monthly temperature, precipitation accumulation, and solar radiation spanning 1961–2000 were acquired from the WorldClim dataset at 1 km^2^ (30 arc sec) spatial resolution (https://www.worldclim.org, accessed January 20, 2025).

In the case of the cropland frequency features, we followed the approach introduced by [61] and described in the previous sections.

### Soil organic carbon modeling

An RF model was trained using 72 soil, climatic, and agronomic features (see S2 Table). A variable-importance analysis was used to identify the most influential features impacting SOC. Following [62], recursive feature elimination [63] was used to select essential predictive features of SOC. This is an iterative process that ranks significant predictors and removes nonexplanatory ones. The process continues until a convergence criterion is met, using the coefficient of determination (*R*^2^) as the evaluation metric.

Additionally, a sensitivity analysis was conducted by grouping rotation features, climatology-based features, and soil properties separately to assess their influence on SOC prediction. Four scenarios were explored: (i) including all 72 features as a benchmark (Case 1); (ii) removing climatology, yielding 24 features (Case 2); (iii) omitting crop rotation history, leaving 64 features (Case 3); and (iv) excluding soil properties, resulting in 57 features (Case 4).

Model performance was assessed using *k*-fold cross-validation, where the dataset was split into 5 folds, each fold used once for validation while the remaining folds were used for training. The *R*^2^ and root mean square error (RMSE) were used to evaluate the ability of the ML model to predict observed SOC. *R*^2^ represents the proportion of variance explained by the model, and RMSE indicates the accuracy of the predicted values [64]. *R*^2^ and RMSE were calculated as follows:

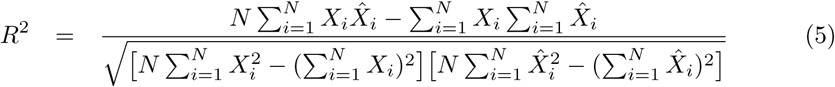

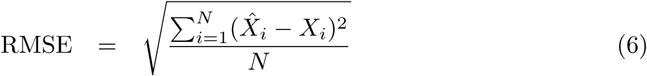

where ^*X_i_*, *X_i_*, and *N* are the predicted SOC, observed SOC, and total number of SOC sample values, respectively. A higher *R*^2^ (close to 1) and lower RMSE (close to 0) indicate better model performance.

Partial Dependence Plots (PDP) were used to visualize the relationship between features and SOC predictions, showing whether the relationship is linear, monotonic, or complex. For this purpose, the implementation available in the pdp library [65] was used within the R environment [40].

## Results

### Phenological-based multi-year cropland classification model

A total of 788 observations were analyzed, with 49.7%, 13.8%, 10.1%, 7.9%, 6.9%, 6.3%, and 5.2% representing potato, fallow, barley, oat, beans, pasture, and lupin, respectively (see Fig. 2A). Most fields are small, clustered below 0.5 ha, with a strongly right-skewed distribution, indicating that while few larger fields exist, they are rare, reaching up to about 8 ha (Fig. 2B).

**Fig 2.**
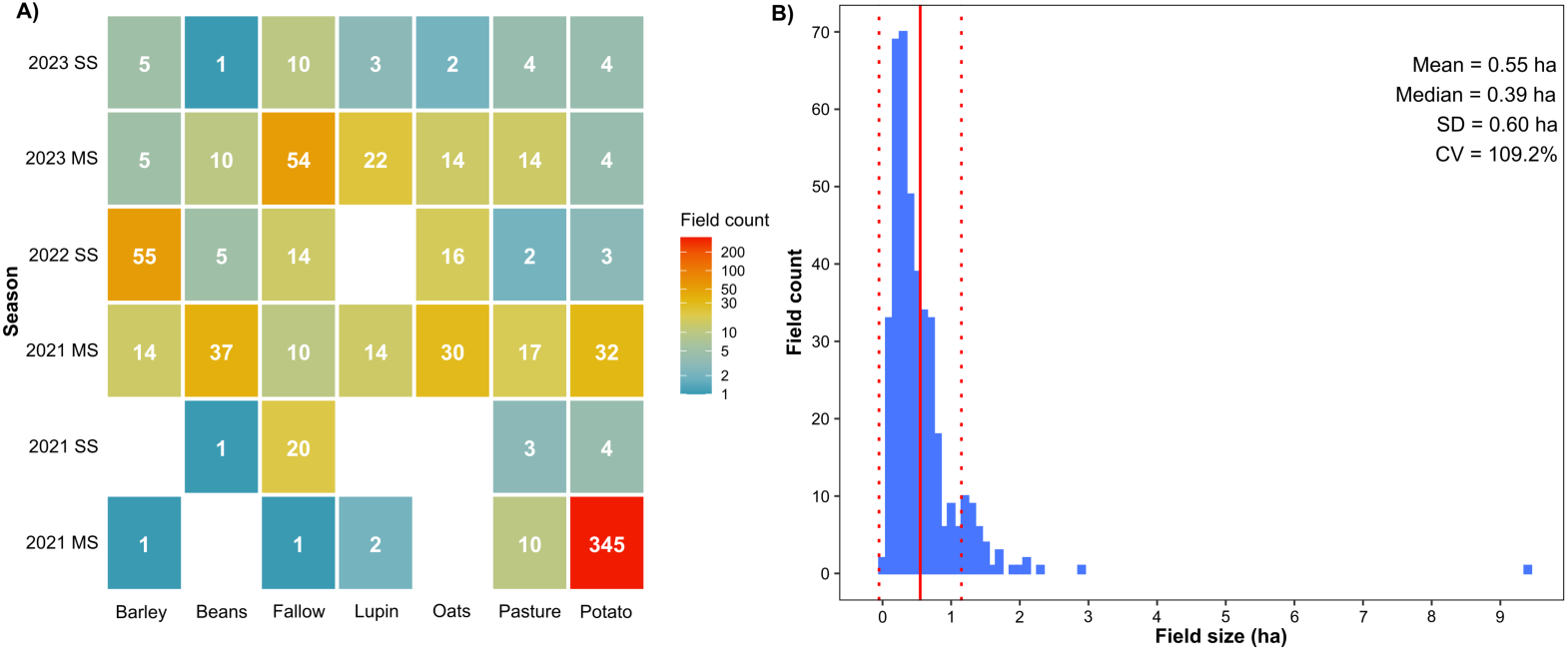
Characterization of farmers’ parcels assessed in the study area. (A) Number of fields for each cropland type per crop season recorded via field farmers’ visits and surveys—main season (MS) and secondary season (SS). (B) Field size distribution of farmers’ fields where SOC was measured.

The adjustment of the Gaussian function showed an average *R*^2^ of 0.62 ± 0.17. The best average performance showed the following pattern by cropland: beans (0.64 ± 0.18) *>* oats (0.63 ± 0.16) *>* barley (0.62 ± 0.18) *>* potato (0.60 ± 0.16) *>* fallow (0.59 ± 0.15) *>* tarwi (0.56 ± 0.15) *>* pasture (0.55 ± 0.15) (Fig. 3). Models fitted better in the secondary campaign for most crops (except for lupin) than the main campaign, with a global *R*^2^ average of 0.65 ± 0.18 and 0.61 ± 0.16, respectively (Fig. 3).

**Fig 3.**
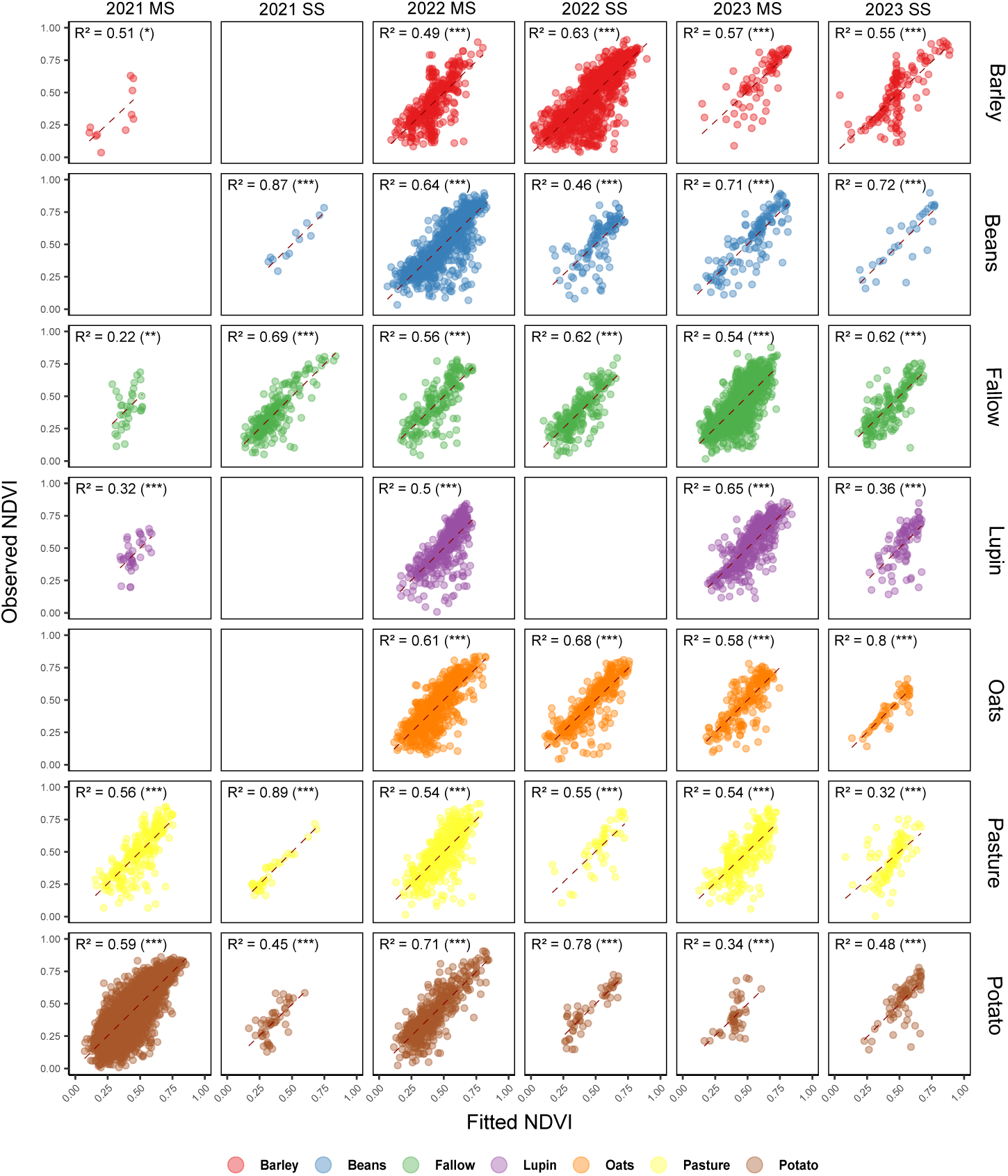
Observed and predicted NDVI using the Gaussian function. Based on the coefficient of Fig(*R*^2^) by crop in the main (MS) and secondary (SS) seasons, 2021–2023. Significance: ns (*p* ≥ 0.05); * (0.05 *> p* ≥ 0.01); ** (0.01 *> p* ≥ 0.001); *** (*p <* 0.001). Symbols indicate statistical significance levels.

NDVI_MAX_ values ranged between 0.55 and 0.80, beans (0.66 ± 0.09) and fallow (0.63 ± 0.09) showed the highest and lowest global average value in this parameter (S2 Fig.), with significant differences among beans vs. fallow–potato, and fallow vs. barley (S3 Table). *µ* values ranged between 4 and 6.5; beans (5.3 ± 1.2) and barley (4.8 ± 1.52) showed the highest and lowest global average value (S2 Fig.), with significant differences among barley vs. all croplands (S3 Table). *σ* values ranged between 3.2 and 4.6; pasture (4.1 ± 0.52) and barley (3.7 ± 0.52) showed the highest and lowest global average value (S2 Fig.), with significant differences among barley vs. all croplands and pasture vs. potato (S3 Table). Crops showed different patterns of clustering using DTW on S2 NDVI-based time series (Fig. 4), with clusters 2, 3, 6, and 7 being the most important in crop differentiation.

**Fig 4.**
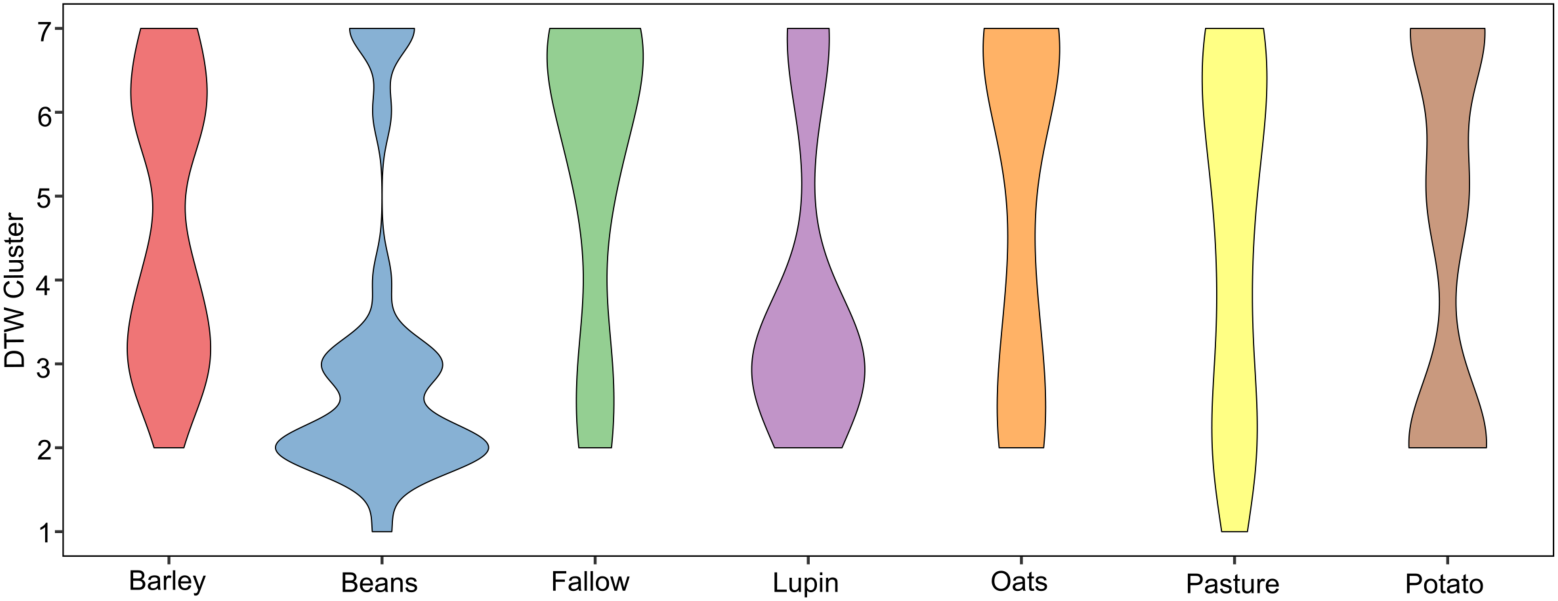
Classification of NDVI time series using Dynamic Time Warping (DTW). The width of the violin diagrams represents the contribution of the corresponding cluster for the characterization of each crop.

The optimal RF architecture for the multi-year cropland classification model was found when mtry = 5, min.node.size = 2, and sample.fraction = 0.88. Validation on the test sample set (30% of the full dataset, *n* = 788 instances) shows strong overall model performance, with F1 scores ranging from 0.81 to 0.98 for the cropland classes (S3 Fig.). The model performance ranking was: lupin (0.98) *>* pasture (0.96) *>* oat (0.93) *>* barley (0.92) *>* fallow (0.91) *>* beans (0.90) *>* potato (0.81). The overall accuracy and Kappa Score were 0.91 and 0.90, respectively (S3 Fig.).

### Soil, Climate, and Crop Rotation features for SOC prediction

SOC values ranged from 2.4 to 22.6 %, with a mean value of 10.61 ± 2.95 % (Table 1). Soils were predominantly loam (34.9 ± 6.88%, 49.4 ± 6.43%, and 24.16 ± 7.70% of Sa, Si, and Cy, respectively), with a moderately acidic pH (4.68 ± 0.85), relatively low EC (0.20 ± 0.24 dS m*^−^*^1^), and moderate to high CEC (15.54 ± 6.81 cmol*_c_* kg*^−^*^1^) (Table 1).

**Table 1.**
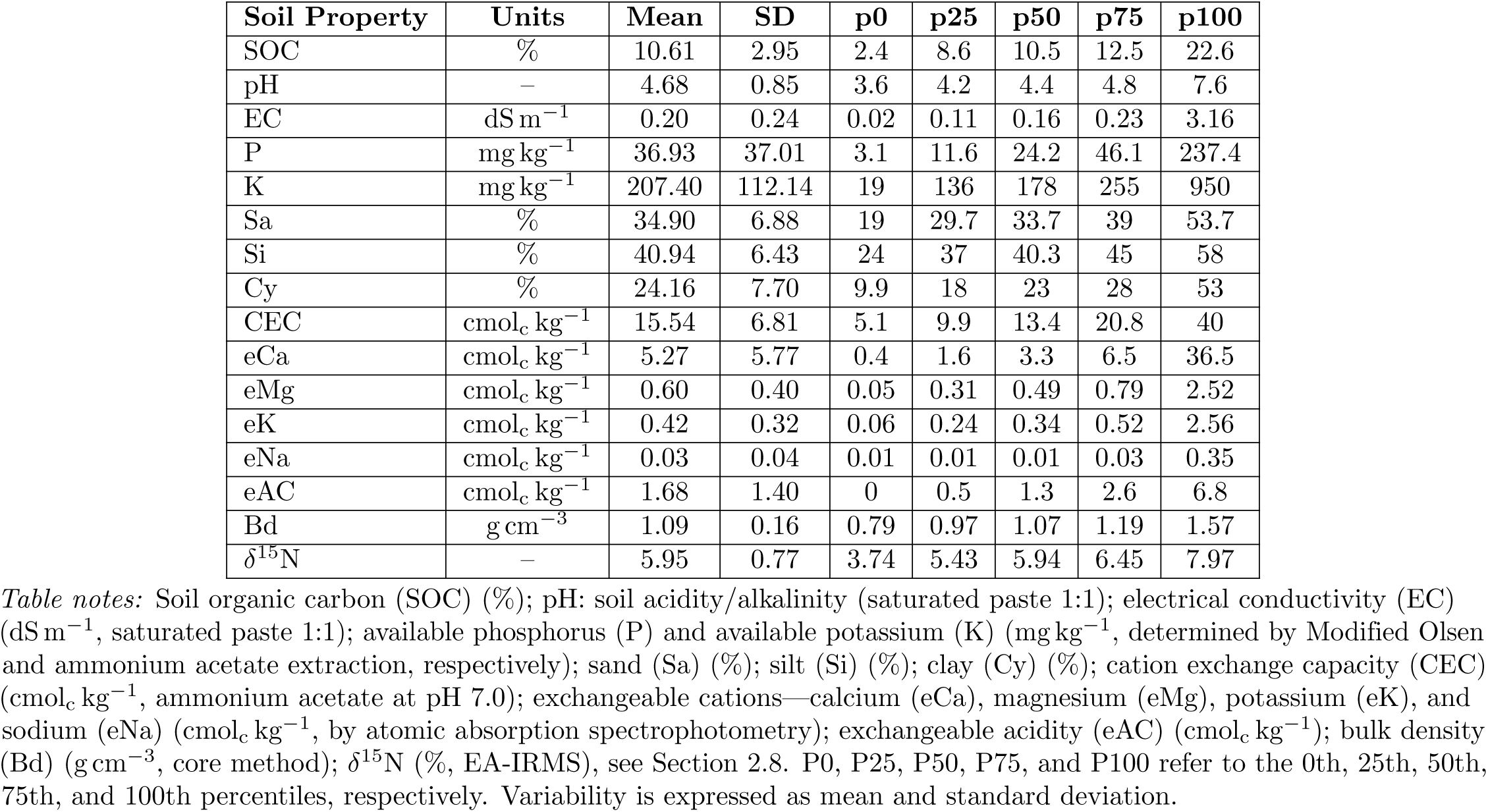
Physical and chemical soil properties.

| Soil Property | Units | Mean | SD | p0 | p25 | p50 | p75 | p100 |
| --- | --- | --- | --- | --- | --- | --- | --- | --- |
| SOC | % | 10.61 | 2.95 | 2.4 | 8.6 | 10.5 | 12.5 | 22.6 |
| pH | – | 4.68 | 0.85 | 3.6 | 4.2 | 4.4 | 4.8 | 7.6 |
| EC | dS m<sup>-1</sup> | 0.20 | 0.24 | 0.02 | 0.11 | 0.16 | 0.23 | 3.16 |
| P | mg kg<sup>-1</sup> | 36.93 | 37.01 | 3.1 | 11.6 | 24.2 | 46.1 | 237.4 |
| K | mg kg<sup>-1</sup> | 207.40 | 112.14 | 19 | 136 | 178 | 255 | 950 |
| Sa | % | 34.90 | 6.88 | 19 | 29.7 | 33.7 | 39 | 53.7 |
| Si | % | 40.94 | 6.43 | 24 | 37 | 40.3 | 45 | 58 |
| Cy | % | 24.16 | 7.70 | 9.9 | 18 | 23 | 28 | 53 |
| CEC | cmol<sub>c</sub> kg<sup>-1</sup> | 15.54 | 6.81 | 5.1 | 9.9 | 13.4 | 20.8 | 40 |
| eCa | cmol<sub>c</sub> kg<sup>-1</sup> | 5.27 | 5.77 | 0.4 | 1.6 | 3.3 | 6.5 | 36.5 |
| eMg | cmol<sub>c</sub> kg<sup>-1</sup> | 0.60 | 0.40 | 0.05 | 0.31 | 0.49 | 0.79 | 2.52 |
| eK | cmol<sub>c</sub> kg<sup>-1</sup> | 0.42 | 0.32 | 0.06 | 0.24 | 0.34 | 0.52 | 2.56 |
| eNa | cmol<sub>c</sub> kg<sup>-1</sup> | 0.03 | 0.04 | 0.01 | 0.01 | 0.01 | 0.03 | 0.35 |
| eAC | cmol<sub>c</sub> kg<sup>-1</sup> | 1.68 | 1.40 | 0 | 0.5 | 1.3 | 2.6 | 6.8 |
| Bd | g cm<sup>-3</sup> | 1.09 | 0.16 | 0.79 | 0.97 | 1.07 | 1.19 | 1.57 |
| δ<sup>15</sup>N | – | 5.95 | 0.77 | 3.74 | 5.43 | 5.94 | 6.45 | 7.97 |

*Table notes:* Soil organic carbon (SOC) (%); pH: soil acidity/alkalinity (saturated paste 1:1); electrical conductivity (EC) (dS m<sup>-1</sup>, saturated paste 1:1); available phosphorus (P) and available potassium (K) (mg kg<sup>-1</sup>, determined by Modified Olsen and ammonium acetate extraction, respectively); sand (Sa) (%); silt (Si) (%); clay (Cy) (%); cation exchange capacity (CEC) (cmol<sub>c</sub> kg<sup>-1</sup>, ammonium acetate at pH 7.0); exchangeable cations—calcium (eCa), magnesium (eMg), potassium (eK), and sodium (eNa) (cmol<sub>c</sub> kg<sup>-1</sup>, by atomic absorption spectrophotometry); exchangeable acidity (eAC) (cmol<sub>c</sub> kg<sup>-1</sup>); bulk density (Bd) (g cm<sup>-3</sup>, core method); δ<sup>15</sup>N (‰, EA-IRMS), see Section 2.8. P0, P25, P50, P75, and P100 refer to the 0th, 25th, 50th, 75th, and 100th percentiles, respectively. Variability is expressed as mean and standard deviation.

Available P and K showed high variability, ranging from 3.1 to 237.4 mg kg*^−^*^1^ and from 19.0 to 950.0 mg kg*^−^*^1^, respectively (Table 1). Mean values of exchangeable bases were 5.27 ± 5.77, 0.60 ± 0.40, 0.42 ± 0.32, and 0.03 ± 0.04 cmol*_c_* kg*^−^*^1^ for eCa, eMg, eK, and eNa, respectively, while eAC averaged 1.68 ± 1.40 cmol*_c_* kg*^−^*^1^ (Table 1). Mean Bd and *δ*^15^N were 1.09 ± 0.16 g cm*^−^*^3^ (range: 0.79–1.57 g cm*^−^*^3^) and 5.95 ± 0.77 ‰ (range: 3.74–7.97 ‰), respectively (Table 1).

Based on historical climate data, monthly averages varied as follows: air temperature ranging from 3.5–16.9 °C, solar radiation from 13,400–15,580 kJ m*^−^*^2^ day*^−^*^1^, and precipitation from 22.9–154 mm. Regarding CR features, the cropland cover frequencies from 2019 to 2022 showed the following distribution: barley (38.7%) *>* potato (21.8%) *>* beans (12.8%) *>* pasture (8.6%) *>* fallow (6.6%) *>* oats (5.9%) *>* lupin (5.7%) (Fig. 5). Barley was the most dominant crop, covering over 50% of seasons in 73 fields, while potato and beans only dominated in 5 and 2 fields, respectively, and the remaining cropland types rarely dominated any parcel. The least persistent crops, appearing in fewer than 10% of seasons in most fields, were sorted in descending order as follows: oats, fallow, lupin, pasture, and beans.

**Fig 5.**
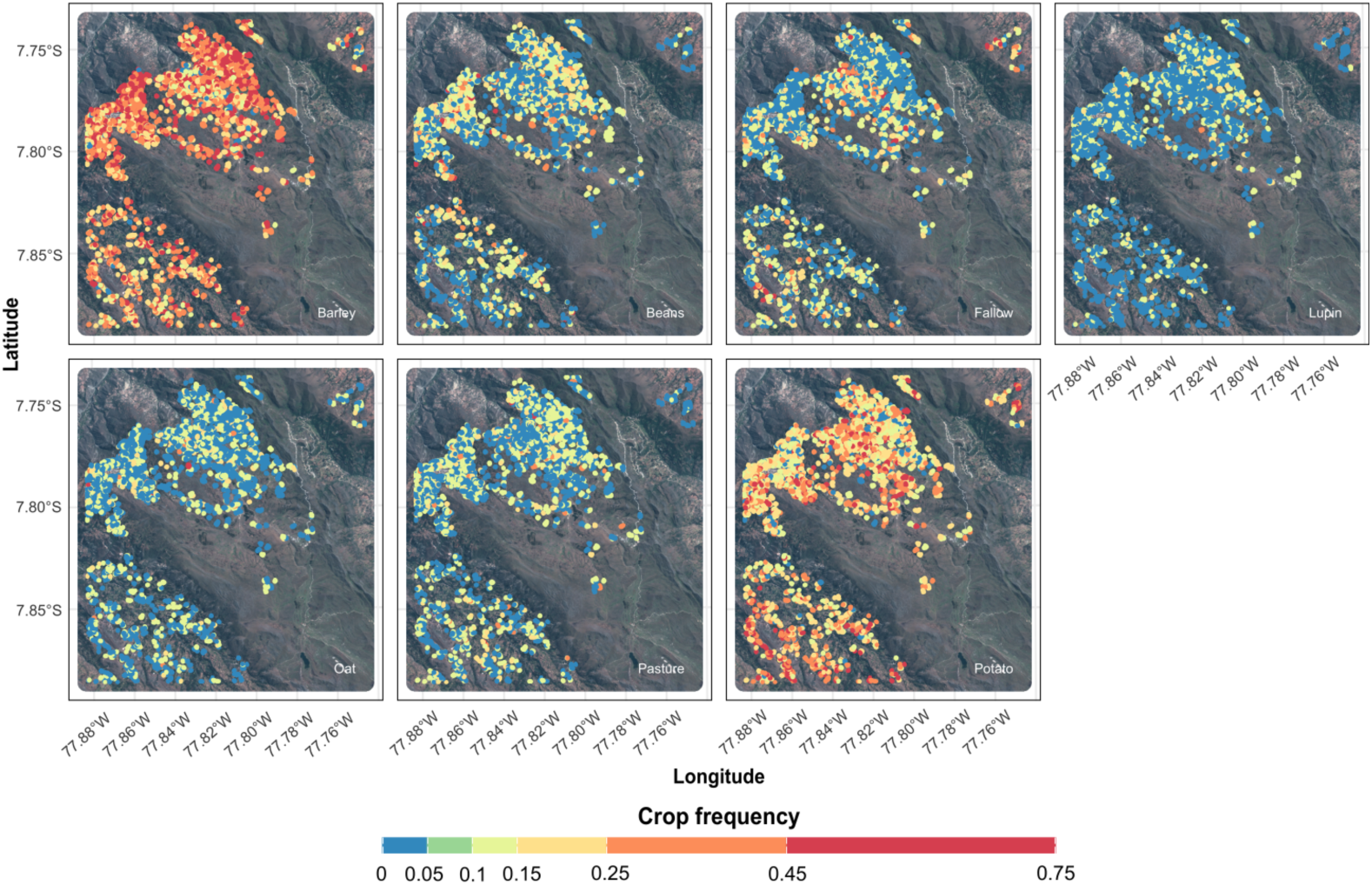
Crop rotation features. Spatial distribution of the features generated by the multi-year cropland classification model from 2019 to 2022, including both surveyed and digitized fields for the study area croplands.

Among the four feature-set configurations, the SOC predictive model performed best in Cases 1 and 3, as evidenced by higher R² values (0.63 in both cases) and lower RMSE values (1.99% and 1.98%, respectively). Conversely, Cases 2 and 4 exhibited comparatively poorer performance, with lower R² values (0.53 and 0.52, respectively) and higher RMSE values (2.24% and 2.28%, respectively) (Fig. 6).

**Fig 6.**
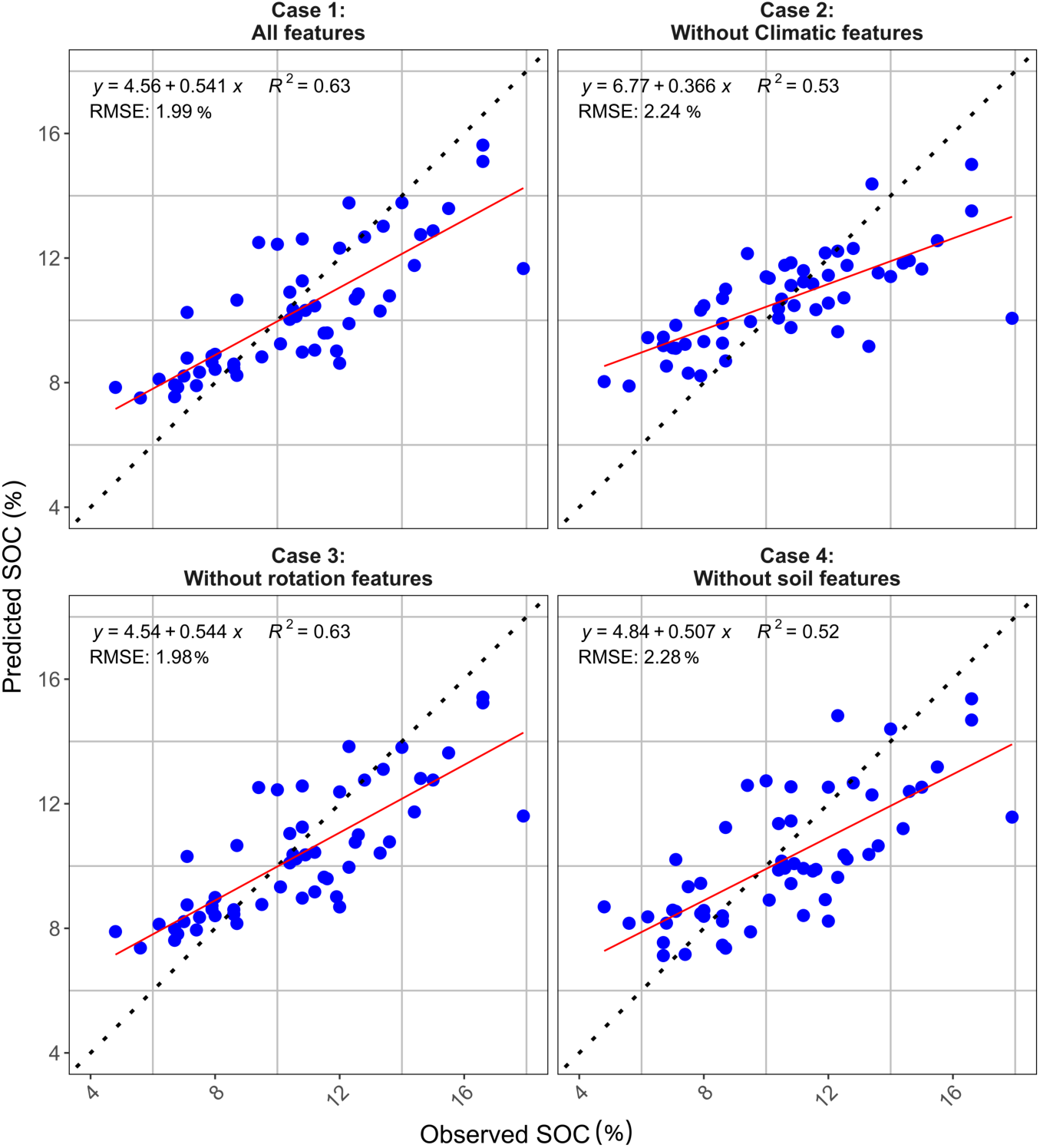
Observed and predicted soil organic carbon (SOC, %). SOC values for four modeling cases: (1) all features included; (2) without climatic features; (3) without rotation features; and (4) without soil features. Solid red lines represent linear regressions between observed and predicted SOC. The dashed line indicates the 1:1 relationship. The regression equation, adjusted determination coefficient (R²), and Root Mean Square Error (RMSE) are shown for each case.

In general, the SOC predictive model showed a tendency toward overestimation for values below ∼10%. In contrast, for values above it, the model showed a tendency toward underestimation (Fig. 6). In Cases 1–3, which included soil features, these variables were among the most important predictors of SOC, with Cy being the most relevant and, together with CEC, EC, eAC, eCa, and eK, consistently ranking among the top 5–7 features (Fig. 7). In contrast, soil pH and the other textural fractions (Sa and Si) showed much lower importance, falling below the top 15 in Case 2 and below the top 20 in Cases 1 and 3 (Fig. 7). In general, CR features ranked higher than climatic features, with fallow frequency being the most critical CR feature (Fig. 7). It ranked first in Case 4 (absence of soil features), while in Case 1 (together with bean frequency) and Case 2, it ranked within the top 9 and 6 respectively (Fig. 7). Potato frequency ranked within the top 20 only in Case 2 (Fig. 7). Among the climatic features, precipitation variables consistently ranked higher than solar radiation.

**Fig 7.**
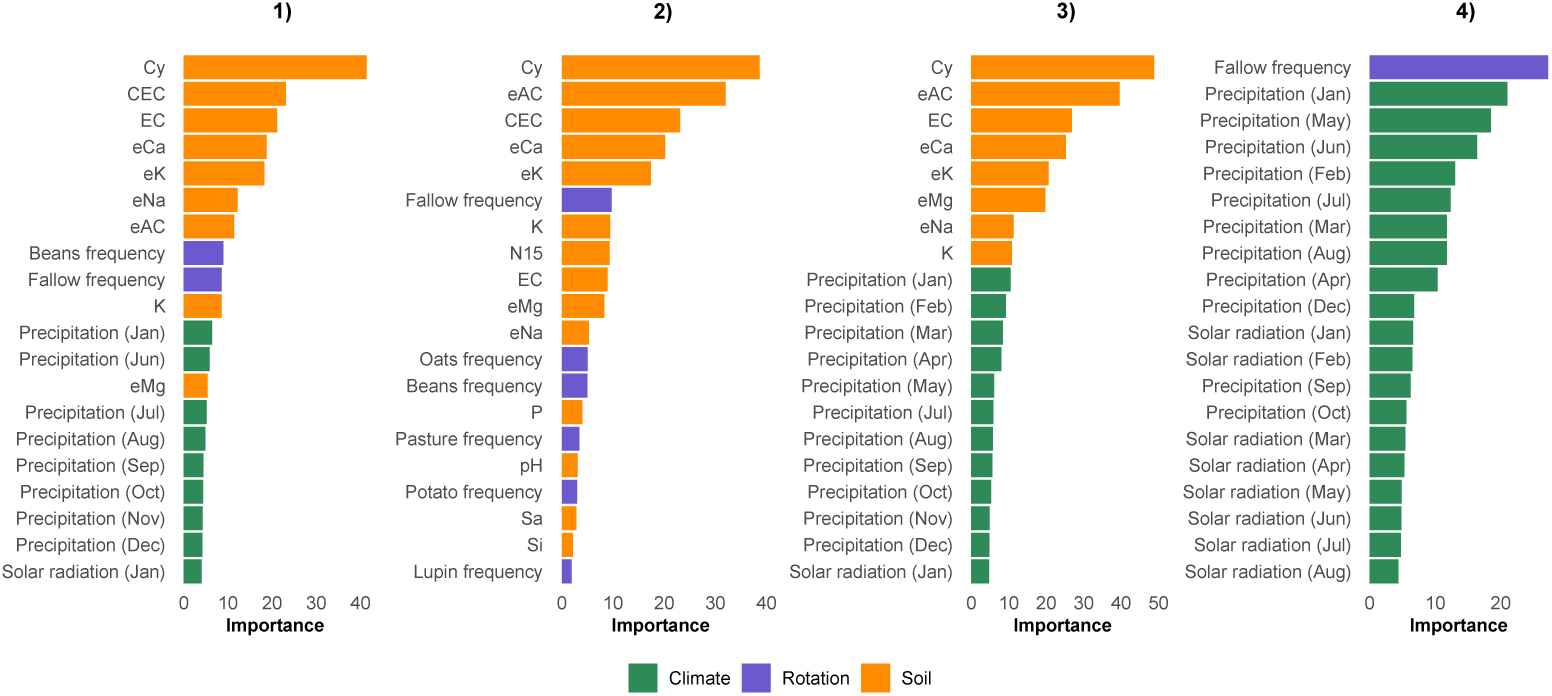
Feature importance of the top 20 variables for predicting SOC. Comparison across four modeling scenarios: Case 1 (all features), Case 2 (without climatic features), Case 3 (without rotation features), and Case 4 (without soil features). Variables are color-coded as green (climate features), purple (rotation features), and orange (soil features). The bars represent the normalized importance of each variable in the model. The soil properties feature abbreviations are described in S2 Table.

Notably, the average precipitation in January was the most significant, ranking within the top 11, 9, and 2 in Cases 1, 3, and 4, respectively. In contrast, temperature variables (both mean and maximum) did not appear among the top 20 in any case (Fig. 7). In Case 4, 19 of the top 20 features corresponded to climatic features (Fig. 7).

The PDP analysis revealed an inverse relationship between Cy and predicted SOC, with sensitivity to SOC variations declining above 20% Cy in Cases 1 and 3, and above 40% Cy in Case 2 (Fig. 8). In contrast, CEC, EC, and eAC exhibited a direct proportional relationship with predicted SOC (Fig. 8). SOC changes in response to CEC were notably faster below 12 cmol*_c_* kg*^−^*^1^ than at higher values in Case 1 and Case 2. In contrast, SOC responded more slowly to eAC in Case 3 compared to Case 2 (Fig. 8). Fallow frequency exhibited a “V”-shaped pattern with predicted SOC, showing an inverse relationship below 25% and a direct relationship above 25% in Case 4 (Fig. 8). Monthly precipitation (January and May) showed a direct relationship with predicted SOC across specific ranges, with minimal sensitivity observed below 115 mm for January and below 40 mm or above 49 mm for May in Case 4 (Fig. 8).

**Fig 8.**
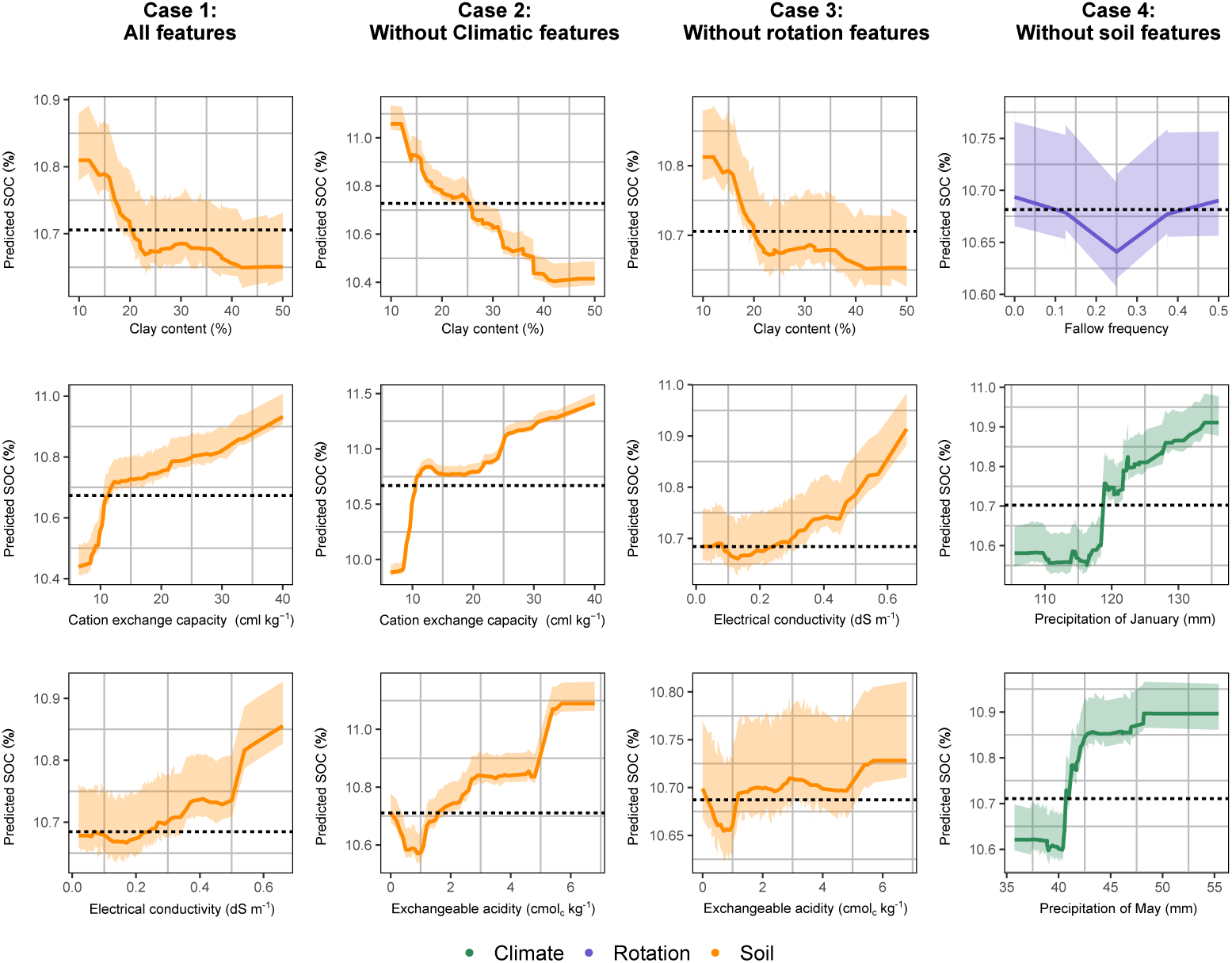
Partial dependent plots (PDP) of predicted soil organic carbon (SOC, %). SOC as a function of top three most important variables for each case: clay content (%), cation exchange capacity (cmol*_c_* kg*^−^*^1^), electrical conductivity (dS m*^−^*^1^), exchangeable acidity (cmol*_c_* kg*^−^*^1^), fallow frequency, and monthly precipitation in January and May (mm). Shaded areas show confidence intervals. Line and ribbon colors indicate the feature category: orange for soil variables, purple for rotation variables, and green for climatic variables. The dashed line represents the mean of observed SOC values.

## Discussion

### Phenology-informed features enable accurate crop classification in heterogeneous Andean smallholder system

In this study, phenological trajectories were modeled with a simple symmetric Gaussian function, parameterized by planting and harvesting dates obtained from field surveys. Despite its parsimony, it requires only three coefficients compared to more complex functions such as the asymmetric Gaussian (four parameters; [66]) and the Double Logistic (five parameters; [67]). The range of *R*^2^ values between observed and fitted NDVI across crops and seasons obtained in this study (0.22–0.89, Fig. 3) reached levels comparable to Gaussian-based analyses of MODIS-EVI time series in boreal forests (0.89–0.94; [68]) and with barley classification in Australia (0.91–0.96; [39]).

Performance was particularly enhanced during the secondary growing season, when reduced rainfall and cloud cover improved the quality of Sentinel-2 imagery (Fig. 1C). Among the extracted parameters, the Gaussian mean (*µ*) proved especially informative, as it reflects the duration of the crop growth cycle. Significant differences in *µ* between barley and other crops (Table S3) underscore its value for distinguishing species with contrasting phenological patterns. NDVI evaluated at *µ* also emerged as a critical indicator of biomass and vigor, consistent with prior research demonstrating its utility for yield prediction in wheat, rice, and cotton [69–71].

The RF classification model achieved strong overall performance, with an accuracy of 0.91 and a Kappa coefficient of 0.90 across seven crop categories. Similarly, RF has proved highly competitive against more complex deep learning models (i.e., CLSTM, 3D U-Net) simulating cloud-prone croplands belonging to smallholders in Africa, particularly when labeled data were limited [13]. This resonates with the study area context, where frequent cloud cover poses similar challenges; yet, RF demonstrated robustness and adaptability across diverse crop categories, even with scarce labeled field data (F1 of 0.71 to 0.93), underscoring the adaptability of time series approaches in smallholder contexts. Nonetheless, classification accuracy varied by crop, reflecting differences in spectral distinctiveness and exposure to seasonal constraints. Lupin and pasture (F1 = 0.98 and 0.96, respectively) yielded the highest accuracies, confirming the ability of Sentinel-2 time series to capture strongly contrasting growth patterns. The performance achieved for lupin is particularly noteworthy, as this study, to our knowledge, represents one of the first documented cases of its reliable discrimination from spaceborne optical imagery. Similarly, the accuracy of pasture classification aligns with studies that successfully integrated Sentinel-1 and -2 data [72] or multi-level neural network approaches [73], reinforcing the conclusion that temporal optical data are sufficient for identifying this crop type.

Other categories also displayed high performance: oat (0.93), barley (0.92), and fallow (0.91). These findings echo recent evidence demonstrating the separability of these classes using temporal metrics. For instance, oats have been reliably identified at the parcel level using Sentinel-2 imagery [74, 75], while barley has been differentiated from wheat using spectral features and classical classifiers [76]. Likewise, fallow land has been effectively mapped using multi-sensor time series and phenological curve fitting, with F1 scores above 0.95 [77]. That such accuracies were obtained in the present study, using a considerably smaller training dataset (224 ha versus more than 2000 ha in comparable work), underscores the robustness of the approach. By contrast, beans (0.90) and potatoes (0.81) exhibited lower classification accuracies. This decline is consistent with previous findings that highlight challenges in separating beans from other legumes and cereals [78]. Potato discrimination is minimal when cultivation coincides with the rainy season (Fig. 1C), as cloud cover and incomplete time series reduce the availability of reliable spectral data [79]. These limitations highlight that classification accuracy can be affected when crop cycles overlap with challenging atmospheric conditions, even with robust phenology-based approaches, and to some extent because of survey data uncertainty.

### In the absence of comprehensive soil measurements, crop rotations provide an effective alternative for modeling SOC

Clay content exhibited a robust and inverse relationship with SOC. Partial dependence analysis revealed that SOC declined sharply in soils with clay contents exceeding 20%, while consistently lower SOC levels were also observed when the CEC fell below 10 cmolc kg*^−^*^1^. These findings align with global reviews showing that clay is often the most influential soil predictor of SOC [8] and digital soil mapping studies where gridded clay, sand, and silt covariates explained significant variation in SOC [80, 81].

Nevertheless, the reported importance of clay varies across contexts. For instance, [82] found climate covariates outweighed soil predictors in Argentina, while [83] identified soil depth as the dominant feature in Canada. Such divergences reflect scale, data resolution, and environmental conditions. Since global products like SoilGrids interpolate across broad regions [84, 85], climatic factors often overshadow soil variables [86]. By contrast, local studies, as in the present work, highlight the strong explanatory role of clay. Mechanistically, clay influences SOC through surface adsorption and aggregate occlusion, protecting organic matter from microbial decomposition [87, 88]. However, accumulation also depends on land use and climate [89–91]. Thus, while clay concentration is consistently important, its role is modulated by context, reinforcing the need for site-specific analyses.

Although soil features were dominant, fallow frequency emerged as the most critical predictor of SOC in crop rotations. When soil data were excluded, fallow frequency, combined with climatic features, explained over half of the SOC variability. Moderate frequencies of fallow were associated with higher SOC, consistent with studies showing that rotation history enhances soil health [57, 61, 92, 93]. Long cropping frequencies [94], and the reduction of soil disturbances through permanent crop cover, rather than annual crops [86], has a positive impact on SOC levels. These findings emphasize that CR data, readily accessible through remote sensing, can serve as practical surrogates for costly soil sampling, enabling moderate prediction performance without the need for extensive laboratory analyses. Potato frequency was not correlated with SOC, contrary to the expectation that intensive tillage associated with potatoes would result in reduced carbon. Tillage has been shown to accelerate soil carbon losses in conventional systems [95]. However, Andean potato-based systems often involve less soil disturbance [96], which may mitigate these effects. This highlights the need for context-specific assessments of management practices. The traditional Highland Andean agriculture established multiyear fallow periods to ensure soil fertility and the recovery of soil functionality, as a key component of complex CR [17]. However, the increasing food demand from cities and markets has driven a reduction in the length of fallow periods [97]. More broadly, the role of fallow in Andean agriculture remains a complex issue [98].

Thus, in the highlands of the Central Peruvian Andes, where Asian markets promoted the cultivation of maca (*Lepidium meyenii* ), an energetic root, [21] did not find significant differences in SOC among fallows of varying ages after maca cultivation. In this ecosystem, [19] reported higher SOC in maca than in fallow plots caused by the incorporation of manure during the management of this crop. On the other hand, in a recent study, [99] pointed out that the seeding of pastures and legumes of traditional fallows has the potential to increase SOC and soil health regeneration in the long term in the Peruvian Central Andes. Such examples highlight the diversity of fallow practices across the Andes, where crop expansion into grasslands and upward shifts in cultivation [100] disrupt the balance between native ecosystems, rotations, and soil carbon storage. Thus, while fallow frequency emerges as a key predictor, its ecological implications require careful contextualization.

Climatic influences add further complexity. Precipitation has consistently been identified as a dominant feature for SOC modeling [82, 86], and its interaction with CR is nonlinear. [8] found SOC increases under rotations were strongest in regions with intermediate mean annual temperature (8–15 *^◦^*C) and precipitation (600–1000 mm), while SOC losses occurred below 120 mm annual rainfall. In the present study area, where annual precipitation ranges from 900 to 1300 mm, high SOC values were obtained (10.61 ± 2.95 %, Table 1). This suggests that favorable climatic conditions may mask the effects of management, complicating the detection of SOC gains from sustainable practices [101]. Future modeling should therefore explicitly account for the interplay between climate and management to better resolve the contributions of CR features.

### Limitations and implications

Although the Sentinel-2 time series effectively characterizes crop phenology [102], its usability is limited by cloud cover. In Chugay, 69.9% and 29.6% of annual scenes were cloud-covered during the main and secondary seasons, respectively, despite the five-day revisit time. Incorporating all-weather observations from non-optical platforms, such as Synthetic Aperture Radar, could address this limitation [103]. The straightforward accessibility of these data (e.g., via GEE; [104]) would facilitate easy integration.

However, Synthetic Aperture Radar data require proper radiometric slope correction and the provision of invalid data masks over areas affected by layover and shadow, particularly in mountainous Andean regions [105]. Integrating Sentinel-1 and -2 time series shows promise [106], although robust multi-sensor workflows remain complex. Further studies are encouraged to advance a multi-sensor approach while maintaining a physiologically sound foundation. Phenological analysis could benefit from established tools like TIMESAT [107] or R packages such as phenofit [68] rather than custom scripts. Since agricultural management decisions are generally made at the parcel level, object-based, rather than per-pixel analysis, is recommended to reduce pixel noise and limit error propagation [106, 108].

While EO information is currently easily accessible, greatly facilitating local and regional studies on crop mapping for sustainable development [109], the most critical gap for Andean crop mapping is reference data. Unlike other regions [13, 110, 111], the Andes lack comprehensive EO datasets linked with field cropping information. We encourage systematic data collection with quality assessment protocols. Since Andean agriculture is highly complex and parcel sizes are small, grouping similar crops into homogeneous categories can be necessary, as distinguishing between crops with similar spectral signatures or growth cycles is a common source of error. Field monitoring at multiple dates, along with a cropping calendar validated by farmers [112], and strategies for sparse data modeling are essential [49]. Regarding field data quality, methods such as hierarchical clustering or self-organizing maps [113] can help reduce spectral confusion and address data imbalance (e.g., grouping barley and oats).

In the study area, the mean values of SOC measured (10.6%) overpasses the values reported in highland Andean grasslands in Peru (4–6%; [19, 21]). This highlights the importance of modeling SOC under this context to distinguish the crucial factors that affect C cycling in this ecosystem. Accurate prediction of SOC in the Andes requires not only technical advances in remote sensing and modeling but also strong coordination among stakeholders to ensure that field data collection and management practices are both scientifically robust and socially viable. The heterogeneity of mountain landscapes—shaped by steep environmental gradients, diverse management systems, and climate variability—makes SOC modeling particularly prone to uncertainty, underscoring the need for joint sampling protocols, standardized databases, and shared monitoring platforms developed by universities, non-governmental organizations, government agencies, and local communities [114–116]. Such collaborative approaches enable the integration of scientific insights by research institutions on practices like crop rotations and fallow periods, while non-governmental organizations and farmers contribute essential knowledge of local feasibility and cultural acceptance, thereby strengthening the credibility of SOC projections.

At the same time, robust monitoring, reporting, and verification protocols are indispensable to link SOC modeling with emerging carbon market mechanisms, where innovations in digital monitoring, reporting and verification, such as remote sensing, big data, and cloud infrastructures, are improving transparency, reducing transaction costs, and aligning local initiatives with international standards [117, 118]. However, implementing these systems in Andean contexts faces constraints related to technical capacity, costs, and governance of data ownership, which highlights the need for pilot initiatives that couple multi-stakeholder coordination with open-access monitoring, reporting, and verification infrastructures [119]. Such integration not only reinforces the scientific basis of SOC monitoring but also positions Andean communities to participate in future carbon markets with greater equity and long-term sustainability.

## Conclusion

To our knowledge, this study is the first to estimate historical field-scale crop rotations and incorporate them as predictors in SOC modeling within the Andes region. The results demonstrate that phenology-driven classification frameworks can reliably discriminate crops even in heterogeneous Andean agroecosystems characterized by small plots and diverse management practices. The integration of Sentinel-2 time series with phenological feature extraction and machine learning modeling effectively addresses many of these challenges. While studies considering fewer crop categories generally report higher accuracy, the successful classification of seven distinct classes in this study demonstrates both the scalability and robustness of the proposed approach. Beyond crop mapping, such reliable discrimination of crop rotations provides critical inputs for downstream SOC modeling.

Overall, the study reveals a hierarchy of predictors for SOC, with clay content exerting the strongest influence, followed by fallow frequency and precipitation. These results suggest that although soil properties remain indispensable, crop rotation and climate variables provide complementary information—particularly in contexts where soil data are limited or costly to obtain. Notably, fallow detection through remote sensing shows strong potential for improving SOC monitoring in Andean landscapes. Nevertheless, the diversity of fallow practices and the confounding influence of climate underscore the need for locally calibrated models. The integration of soil, rotation, and climate features therefore offers a more nuanced understanding of SOC dynamics, with direct implications for designing sustainable land management strategies in fragile mountain agroecosystems.

## Supporting information

Modeled crop types in the Chugay study area.

Classification features for multi-year cropland classification using Sentinel-2 imagery A and B time series (S2A, S2B).

Boxplots of NDVI-derived phenological parameters.

Feature sets used in the soil organic carbon sensitivity analysis.

Confusion matrix for the test set.

Kruskal-Wallis test and mean rank differences of Dunn&#8217s post hoc test results.

## Supporting information

**S1 Table. Classification Features.** Classification features for multi-year cropland classification using Sentinel-2 imagery A and B time series (S2A, S2B) from the Harmonized Sentinel-2 Multispectral Instrument (MSI). Blue, green, red, near-infrared (NIR), shortwave infrared 1 (SWIR1), and shortwave infrared 2 (SWIR2) are surface reflectance values from Band 2 (blue, 496.6 nm (S2A)/492.1 nm (S2B)), Band 3 (green, 560 nm (S2A)/559 nm (S2B)), Band 4 (red, 664.5 nm (S2A)/665 nm (S2B)), Band 8 (NIR, 835.1 nm (S2A)/833 nm (S2B)), Band 11 (SWIR1, 1613.7 nm (S2A)/1610.4 nm (S2B)), and Band 12 (SWIR2, 2202.4 nm (S2A)/2185.7 nm (S2B)). *f* = 0.05.

**S2 Table. Feature sets used in the soil organic carbon sensitivity analysis.**

**S3 Table. Kruskal–Wallis test and mean rank differences of Dunn’s post hoc test results.** Results with Holm’s correction method comparing Gaussian parameters across different cropland cover categories. NDVI_MAX_ is the Normalized Difference Vegetation Index (NDVI) value at peak canopy cover time, *µ* is the time to reach peak growth, and *σ* is the full width at half maximum, signaling the length of the growing period. The significance codes are: *** for *p <* 0.001, ** for *p <* 0.01, * for *p <* 0.05, · for *p <* 0.1, and no symbol for *p* ≥ 0.1.

**S1 Fig. Modeled crop types in the Chugay study area.** Potato (*Solanum tuberosum* L.) (a), beans (*Vicia faba*) (b), lupin (*Lupinus mutabilis*) (c), fallow (d), pasture (e), barley (*Hordeum vulgare* L.) (f), and oats (*Avena sativa* L.) (g).

**S2 Fig. Boxplots of NDVI-derived phenological parameters.** (A) Maximum Normalized Difference Vegetation Index (NDVI) value at peak canopy cover time (NDVI_MAX_), (B) time to reach peak growth (*µ*), and (C) full width at half maximum (*σ*), estimated from Gaussian functions fitted to Sentinel-2 NDVI time series for each cropland type.

**S3 Fig. Confusion matrix for the test set.** Test set corresponds to 30% of the full dataset (*n* = 788 instances).

## Acknowledgments

We sincerely thank all the farmers from the communities in the Chugay district who generously shared information for this study. We also thank the Google Earth Engine team for providing the free platform for Sentinel-2 data processing.

## Author Contributions

**Conceptualization:** Marcelo Bueno, Hildo Loayza.

**Data curation:** Marcelo Bueno.

**Formal analysis:** Marcelo Bueno, Hildo Loayza.

**Supervision:** Hildo Loayza, David A. Ramírez.

**Writing – original draft:** Marcelo Bueno.

**Writing – review & editing:** Hildo Loayza, David A. Ramírez, Johan Ninanya.

**Methodology:** Marcelo Bueno, Hildo Loayza.

**Resources:** Carlos Mestanza, Percy Briceños, Johan Ninanya, Javier Rinza, Ronal Otiniano.

**Funding acquisition:** Jan Kreuze, David A. Ramírez.

## Notes

https://github.com/kundun14/multi-annual-crop-classification-with-phenological-features

https://doi.org/10.5281/zenodo.17352136

### Competing Interest Statement

The authors have declared no competing interest.

### Funding

- Bill & Melinda Gates Foundation: OPP1070785
- Sustainable Farming CGIAR Science Program
- Deutsche Gesellschaft für Internationale Zusammenarbeit (GIZ): 81275993
- CGIAR Initiative on Excellence in Agronomy
- CGIAR Sustainable Farming Science Program

