## Supplementary figures and images for "Can a history of crop rotations improve the prediction of soil organic carbon in the Andes? integrating machine learning multi-annual crop classification as a proxy of soil management"

### Boxplots of NDVI-derived phenological parameters.

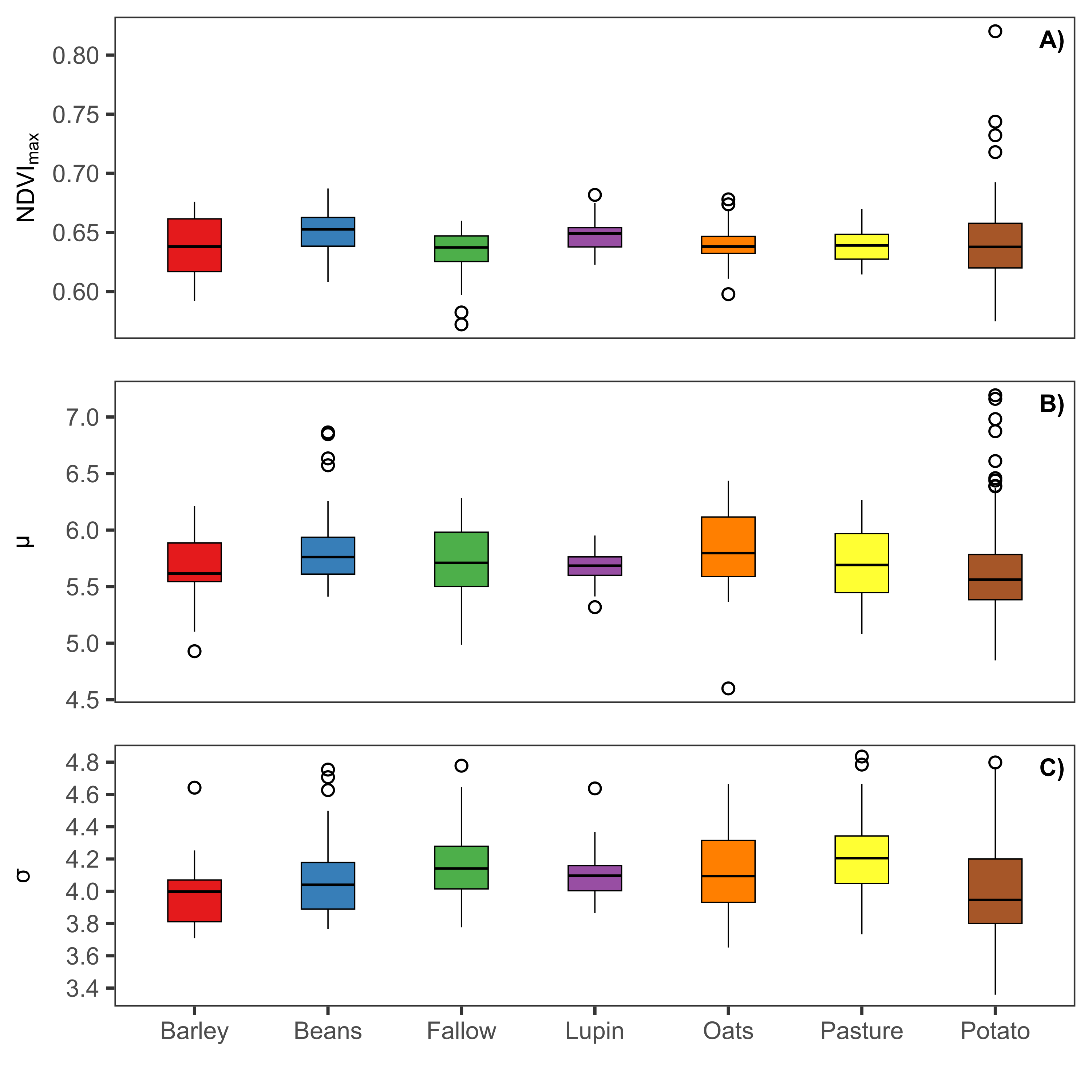

### Confusion matrix for the test set.

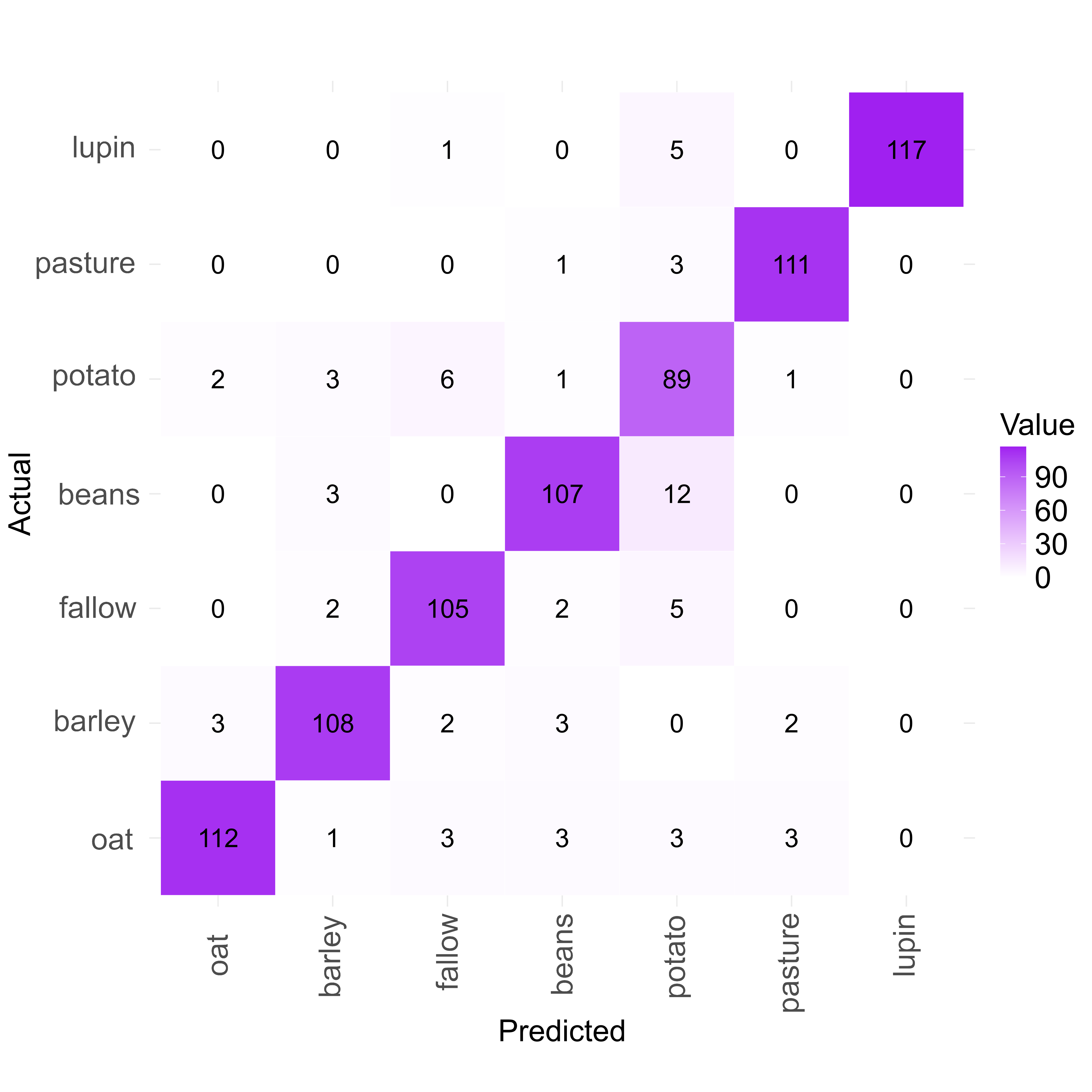

### Modeled crop types in the Chugay study area.

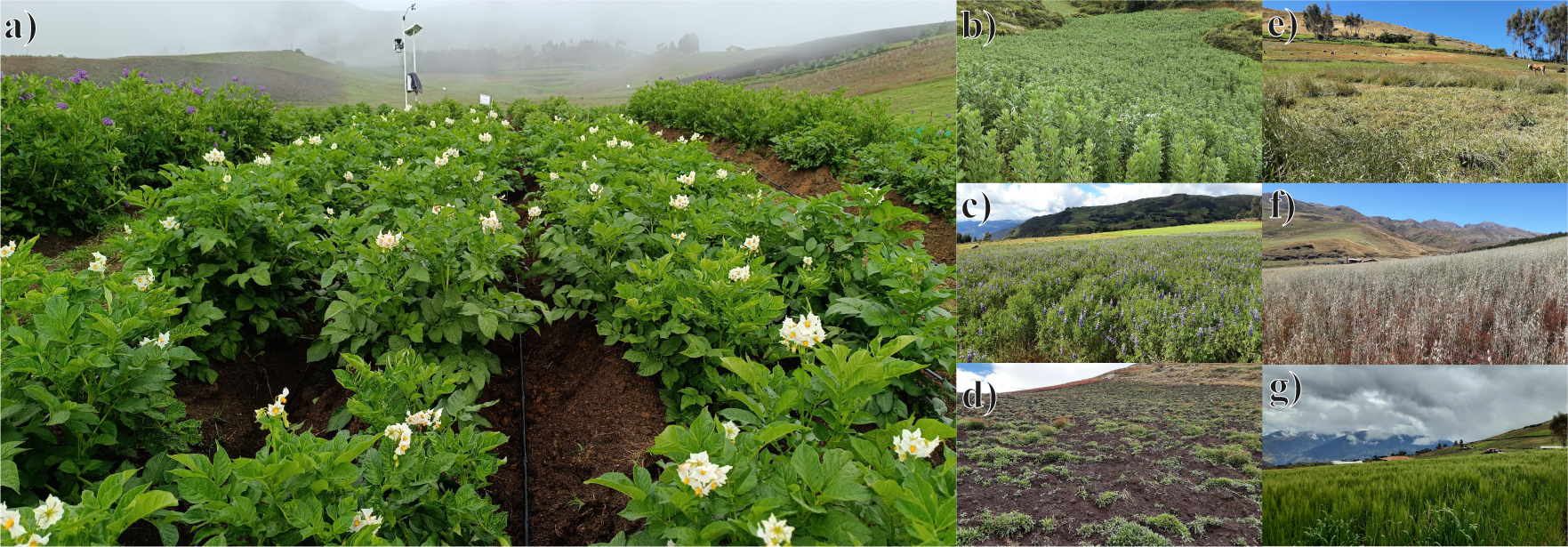
