## Supplementary material for "Can a history of crop rotations improve the prediction of soil organic carbon in the Andes? integrating machine learning multi-annual crop classification as a proxy of soil management": Classification features for multi-year cropland classification using Sentinel-2 imagery A and B time series (S2A, S2B).

|  | Index/Parameter Definition | Acronym/Formula |
| --- | --- | --- |
| Phenology Gaussian Feature | Midpoint of the crop growth curve | µ |
|  | Vegetative period | σ |
| Vegetation Indices | Bare Soil Index (BSI) evaluated at µ | $BSI =\frac{\left( SWIR2+R \right)- (NIR+B)}{\left( SWIR2+R \right) + (NIR+B)}$ |
|  | Normalized Difference Vegetation Index (NDVI) evaluated at µ ($\mathrm{NDVI}_{MAX}$) | $NDVI =\frac{NIR-R}{NIR + R}$ |
|  | Specific Leaf Area Vegetation Index (SLAVI) evaluated at µ | $SLAVI=\frac{NIR}{RED+ SWIR1}+f$ |
|  | Normalized Burn Ratio 2 (NBR2) evaluated at µ | $NBR2= \frac{SWIR1-SWIR2}{SWIR1 + SWIR2}$ |
|  | Normalized Difference Moisture Index (NDMI) evaluated at µ | $\mathrm{NDMI}= \frac{NIR-SWIR2}{NIR+ SWIR2}$ |
| Dynamic Time Warping (DTW) Features | Cropland cluster groups from DTW hard clustering. | DTW_POTATO,_ DTW_LUPIN,_ DTW_FALLOW,_ DTW_BEANS,_ DTW_BARLEY,_ DTW_PASTURE,_ DTW_OATS_ |
|  | Cropland cluster groups from DTW fuzzy clustering | DTW_FUZZ_ |
