## Supplementary material for "Can a history of crop rotations improve the prediction of soil organic carbon in the Andes? integrating machine learning multi-annual crop classification as a proxy of soil management": Feature sets used in the soil organic carbon sensitivity analysis.

| Feature Group | Variable | Method/Source |
| --- | --- | --- |
| *Soil Features* | Soil pH value (acidity/alkalinity) | Soil-Water suspension 1:1 ^[1]^ |
|  | Electrical conductivity of soil (EC, dS/m) | Soil-Water suspension 1:1 ^[2]^ |
|  | Available Phosphorus (P) and Potassium (K) content in soil (mg/kg) | Modified Olsen method (P) and Ammonium acetate extraction (K). |
|  | Soil texture components: Sand (Sa), Silt (Si), and Clay (Cy) fractions (%) | Sedimentation method by Bouyoucus hydrometer. |
|  | Cation exchange capacity (CEC, cmol_c_/kg) | Ammonium acetate buffered at pH 7.0 |
|  | Exchangeable bases: Calcium (eCa), Magnesium (eMg), Potassium (eK), and Sodium (eNa) (cmol_c_/kg) | Atomic absorption spectrophotometry |
|  | Exchangeable acidity (eAC, cmol_c_/kg) | Extraction by 1N KCl and titration with NaOH |
|  | Bulk density (Bd, g/cm³) | Core method |
|  | Nitrogen isotopic ratio (δ^15^N) | Isotope ratio mass spectrometry (EA-IRMS) |
| *Crop Rotation Features* | Frequency of occurrences (F) for respective crops in the dataset | Derived from cropland data |
| *Climatology Features* | Mean monthly precipitation for January (P_1_) to December (P_12_) | [https://www.worldclim.org](https://www.worldclim.org/) |
|  | Mean monthly solar radiation (kJ/m²/day) for months January (R_1_) to December (R_12_) | [https://www.worldclim.org](https://www.worldclim.org/) |
|  | Mean monthly temperature (°C) from January (TM_1_) to December (TM_12_) | [https://www.worldclim.org](https://www.worldclim.org/) |
|  | Mean monthly maximum temperature (°C) from January (TX_1_) to December (TX_12_) | [https://www.worldclim.org](https://www.worldclim.org/) |

1. Bazan, T. R. (2017). Manual de procedimientos de los análisis de suelos y agua con fines de riego. https://repositorio.inia.gob.pe/handle/20.500.12955/504
2. Bazan, T. R. (2017). Manual de procedimientos de los análisis de suelos y agua con fines de riego. https://repositorio.inia.gob.pe/handle/20.500.12955/504
