## Supplementary material for "Can a history of crop rotations improve the prediction of soil organic carbon in the Andes? integrating machine learning multi-annual crop classification as a proxy of soil management": Kruskal-Wallis test and mean rank differences of Dunn&#8217s post hoc test results.

| **Pairwise comparisons** | $\boldsymbol{NDVI}_{\boldsymbol{MAX}}$ | **µ** | **σ** |
| --- | --- | --- | --- |
| Beans vs Barley | 30.03 | 159.19 *** | 88.98 ** |
| Fallow vs Barley | -82.52 * | 99.74 *** | 111.63 *** |
| Lupin vs Barley | -2.14 | 143.01 *** | 118.11 *** |
| Oats vs Barley | -44.99 | 98.07 ** | 87.35 * |
| Pasture vs Barley | -50.15 | 114.71 ** | 135.16 *** |
| Potato vs Barley | -47.24 | 99.33 *** | 56.97 * |
| Fallow vs Beans | -112.55 *** | -59.44 | 22.66 |
| Lupin vs Beans | -32.17 | -16.17 | 29.13 |
| Oats vs Beans | -75.01 | -61.12 | -1.63 |
| Pasture vs Beans | -80.18 | -44.47 | 46.18 |
| Potato vs Beans | -77.26 ** | -59.86 | -32 |
| Lupin vs Fallow | 80.38 | 43.27 | 6.47 |
| Oats vs Fallow | 37.53 | -1.68 | -24.29 |
| Pasture vs Fallow | 32.37 | 14.97 | 23.53 |
| Potato vs Fallow | 35.28 | -0.42 | -54.66 |
| Oats vs Lupin | -42.84 | -44.95 | -30.76 |
| Pasture vs Lupin | -48.01 | -28.3 | 17.05 |
| Potato vs Lupin | -45.09 | -43.69 | -61.13 |
| Pasture vs Oats | -5.16 | 16.65 | 47.82 |
| Potato vs Oats | -2.25 | 1.26 | -30.37 |
| Potato vs Pasture | 2.91 | -15.39 | -78.19 * |
